# Phenotypic heterogeneity is adaptive for microbial populations under starvation

**DOI:** 10.1101/2021.04.05.438545

**Authors:** Monika Opalek, Bogna Smug, Michael Doebeli, Dominika Wloch-Salamon

**Affiliations:** Jagiellonian University, Faculty of Biology, Institute of Environmental Sciences, Kraków, Poland; Malopolska Centre of Biotechnology, Jagiellonian University, Kraków, Poland; University of British Columbia, Vancouver, Canada

**Author notes:** Address correspondence to: D. Wloch-Salamon, and, M. Opalek,.

## Abstract

To persist in variable environments populations of microorganisms have to survive periods of starvation and be able to restart cell division in nutrient-rich conditions. Typically, starvation signals initiate a transition to a quiescent state in a fraction of individual cells, while the rest of the cells remain non-quiescent. It is widely believed that, while quiescent cells (Q) help the population to survive long starvation, the non-quiescent cells (NQ) are a side effect of imperfect transition. We analysed regrowth of starved monocultures of Q and NQ cells compared to mixed, heterogeneous cultures in simple and complex starvation environments. Our experiments, as well as mathematical modelling, demonstrate that Q monocultures benefit from better survival during long starvation, and from a shorter lag phase after resupply of rich medium. However, when the starvation period is very short, the NQ monocultures outperform Q and mixed cultures, due to their short lag phase. In addition, only NQ monocultures benefit from complex starvation environments, where nutrient recycling is possible. Our study suggests that phenotypic heterogeneity in starved populations could be a form of bet hedging, which is adaptive when environmental determinants, such as the length of the starvation period, the length of the regrowth phase, and the complexity of the starvation environment vary over time.

**Importance:** Non-genetic cell heterogeneity is present in glucose starved yeast populations in the form of quiescent (Q) and nonquiescent (NQ) phenotypes. There is evidence that Q cells help the population to survive long starvation. However, the role of the NQ cell type is not known, and it has been speculated that the NQ phenotype is just a side effect of imperfect transition to the Q phenotype. Here we show that, in contrast, there are ecological scenarios in which NQ cells perform better than monocultures of Q cells or naturally occuring mixed populations containing both Q and NQ. NQ cells benefit when the starvation period is very short and environmental conditions allow nutrient recycling during starvation. Our experimental and mathematical modeling results suggest a novel hypothesis: the presence of both Q and NQ phenotypes within starved yeast populations may reflect a form of bet hedging, where different phenotypes provide fitness advantages depending on environmental conditions.

## Introduction

### Stationary phase is heterogenous

Survival of microbial populations depends on the individual cells’ ability to adjust their phenotype in response to challenging environmental conditions. Structured environments, ageing, and nutrient limitation have been identified as factors driving non-genetic heterogeneity visible as multiple cellular phenotypes present in microbial populations [1–3]. In particular, transition to quiescent or other spore-like cell type induced by starvation is a phenotype of fundamental importance in medical microbial biology. Not only does it play a crucial role in biofilm forming [4], but it is also significant in cancer formation [5]. Quiescence is well studied in the yeast *Saccharomyces cerevisiae*, where a fraction of cells undergo specific molecular and cellular reprogramming, and actively cease division, when there is a lack of essential resources. As a consequence, in the stationary phase, genetically clonal yeast population contains a mixture of non-quiescent (NQ), quiescent (Q) and dead cells.

The quiescent phenotype is complex, and its precise characterisation is the subject of ongoing research [6–8]. Q cells’ testing is also extremely challenging, because once they re-enter the cell cycle they are no longer in the quiescent state. Yet, studying quiescence in *S. cerevisiae* has the unique advantage of the possibility of obtaining the fractions of Q and NQ cells by centrifugation on a density gradient [9–12]: Q cells are gathered in the denser, lower fraction, while the upper, less dense fraction predominantly consists of NQ cells. Quiescent yeast cells can be characterised by a thickened cell wall, dense vacuole and an accumulation of storage materials such as trehalose [5,9,13,14]. While starved, NQ cells stop at various stages of the mitotic cell cycle and do not undergo transition to Q. As a consequence, NQ cells vary in internal organisation and are more heterogeneous than Q cells. Differentiation into quiescence starts in growing yeast populations after the first signals of starvation, ca. 20 hours of inoculation in glucose rich medium. For common laboratory prototrophic *S*.*cerevisiae* strain S288C Q:NQ cells ratio is about 70:30 [9,11,15,16].

While it is the whole population of Q and NQ cells that experience starvation, Q cells with their adaptation to long starvation survival, stress tolerant viability and higher recovery speed are the ones responsible for population re-growth [9,11,17]. Thus, evolved enrichment of Q cells in starved populations (up to 95%) resulted in significant increase of re-growth abilities after 22 days of starvation [15]. Other research showed that after 4 weeks of starvation, Q cells exhibited 87% viability while only 3% of NQ cells were still viable (counted as CFU [9]). Q cells also survive higher amounts of stress, such as temperature [9,11] and toxins (including antifungals [4]). Regrowth abilities of Q and NQ cells were checked separately after culture fractionations, however, in most of the experimental set ups, stress was applied to the unseparated stationary phase population consisting of certain mixtures of Q and NQ cells [10,18–20].

The Q/NQ population balance can be affected by many factors [17]. It was shown that the Q/NQ ratio can be modified by selection to some extent, however both Q and NQ cells appear to always be present in stationary populations [10,15,21]. This raises the important question of how the Q/NQ balance evolves in various ecological scenarios. Because of the apparently clear advantage of Q cells in stress survival, the presence of NQ cells could be an inevitable by-product of cellular physiology, with no particular adaptive significance. For example, it was hypothesized that replicative ageing is a factor determining the transition to quiescence: the presence of NQ cells in a starved population would simply reflect the inability of old cells to enter quiescence [19]. However recent research using more advanced laboratory techniques questioned this interpretation [11]. Alternatively, the presence of both Q and NQ phenotypes within starved population may reflect some form of bet hedging, where different phenotypes provide fitness advantage depending on environmental conditions [22–24].

### Is stationary phase heterogeneity advantageous for population’s survival?

Here we use population-level experiments to shed light on the adaptive significance of the Q/NQ cell ratio in yeast. We test whether populations composed entirely of Q cells (Q monoculture) have an ecological advantage over natural populations (mix culture with Q and NQ cells in 3:1 ratio), as well as over NQ monocultures. These mixed cultures imitate the naturally occurring Q/NQ cell ratio in starved laboratory S288C strain and were therefore taken as a reference point. We monitored starvation survival of experimental cultures weekly via regrowth experiments (Fig.1., see details in Materials and Methods). We describe populations’ growth curves after various starvation lengths - long starvation scenario lasting from 1 to 6 weeks and short starvation scenario lasting 4 days. We analyse the impact of the environment during long starvation, where cells were suspended either in sterile water (simple environment) or spent medium (complex environment). Finally, we develop a mathematical model to explain our experimental results and to predict ecological outcomes.

Our experiments show that Q monocultures regrow relatively better than mixed cultures if the starvation phase is long enough, and that this advantage diminishes after longer regrowth time. We used the mathematical model to assess possible reasons for this advantage. The model supports the notion that the ecological advantage of Q monocultures is due to lower death rate of Q cells during starvation, and to shorter lag times of Q cell after starvation periods longer than one week. We also demonstrate that, due to nutrient recycling, NQ monocultures do relatively better in complex than simple starvation environment. Finally, we hypothesise based on the model and confirm experimentally that NQ monocultures can regrow faster than Q monocultures when the starvation period is very short. Our conclusion is that the presence of both quiescent and non-quiescent cells could be advantageous for population survival in fluctuating environments because Q cells survive long starvation better and NQ cells restart divisions faster if the starvation period is short. Moreover, when there is a possibility of nutrient recycling in complex starvation environments, NQ cells may suffer less from unfavourable environmental conditions compared to simple environments. This supports the hypothesis that the existence of both cell types in natural populations could be a form of bet-hedging rather than the effect of imperfect transition into quiescence.

## Materials and Methods

### Strain, Q and NQ cell acquisition

We used a derivative of the laboratory haploid *Saccharomyces cerevisiae* strain s288C (Mat α, *ura3::KanMX4*) [Cubillos *et al*. 2009]. Yellow fluorescent marker (YFP) was amplified from genomic DNA from the BY4741-YFP.natR strain and integrated into the ancestral strain according to previously described protocols [26].

In order to obtain Q and NQ cells we applied a previously described fractionation procedure in a density gradient [9]. In short, an overnight culture was diluted tenfold and 100 µl (∼2× 10^7^cells) was incubated on an agar YPD plate for 4 days in 30°C (reaching cells density ∼2×10^8^/ml). The density gradient was obtained by mixing Percoll and NaCl (1,5M) in proportion 9:1 v/v, and by subsequent centrifugation in angular rotor for 20 minutes at rcf = 10078g (centrifuge: MPW Med. Instruments model MPW-352R). Then the culture was washed from the plate (10 ml 50 mM Tris, pH=7.5), and 4 ml was pelleted, placed on the top of a density gradient and centrifuged in swinging-bucket rotor for 60 minutes at rcf = 417g. Upper (NQ cells) and lower (Q cells) fractions were carefully separated by pipets and placed in individual tubes (Fig. 1A). For long starvation scenario, harvested cells were stored at - 70°C in 25% glycerol until the beginning of the starvation experiment. For the short starvation scenario, harvested cells were immediately used to prepare experimental populations and placed for regrowth.

**Fig. 1.**
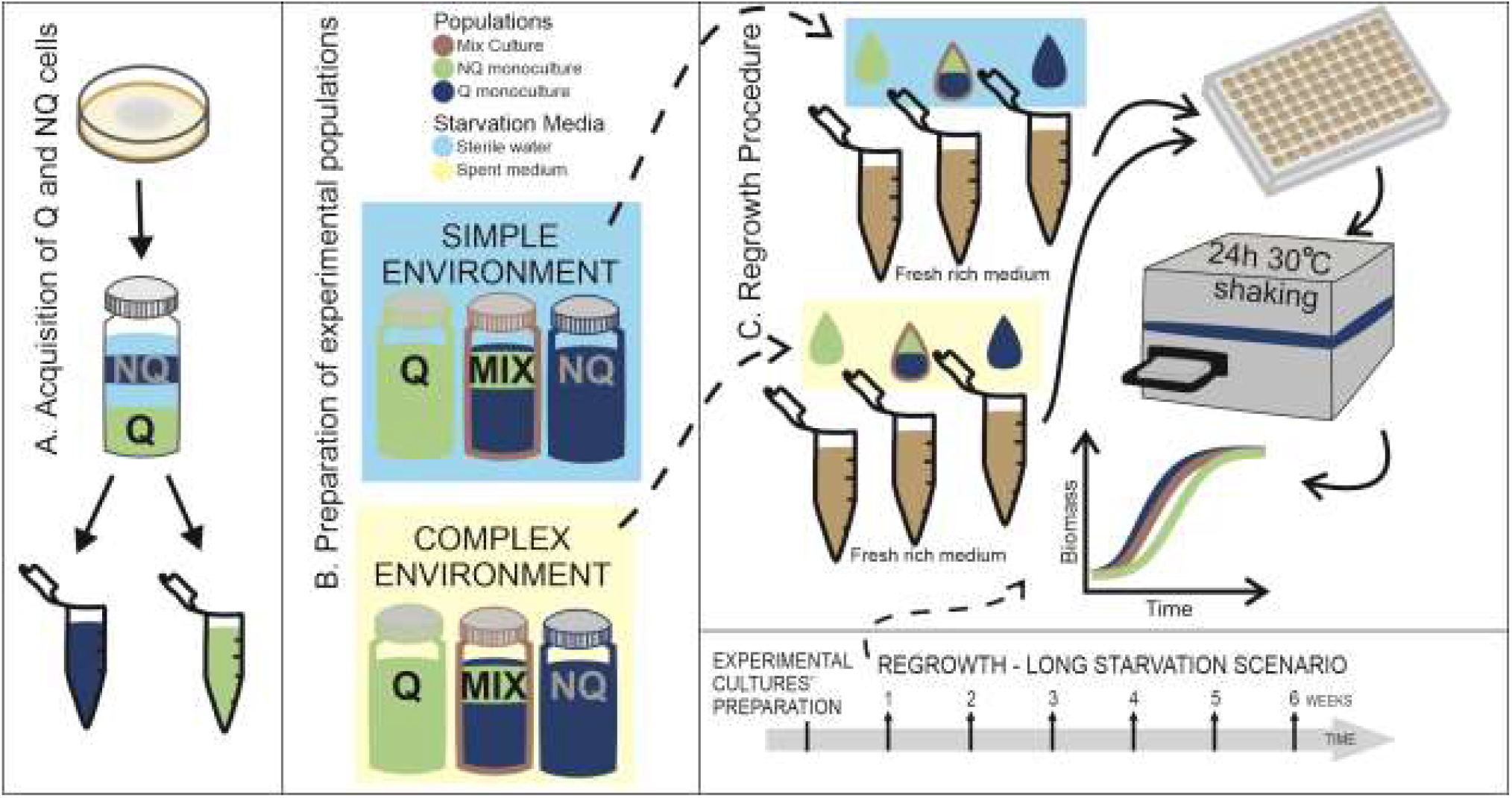
Schematic representation of performed experiments. A – After inoculation, population was growing on agar YPD plate for four days, after that Q (navy) and NQ (green) cells were separated by centrifugation on density gradient. B – Six types of experimental populations were prepared, each type was prepared in five repetitions, giving all together 30 independent cultures. First all Q and NQ cells acquired by fractionation procedure were merged and diluted to equal OD. Next, mix cultures were prepared by mixing Q and NQ cells in 3:1 proportion. Then cells were pelleted and resuspended in starvation medium – sterile water for simple environment (blue background) and spent medium for complex environment (beige background). C – To asses starvation survival, we weekly took samples of experimental cultures. Cells were pelleted and resuspended in fresh rich medium. Then samples were loaded into 96-well plate and placed into plate reader for regrowth. The samples were incubated for 24 hours in 30°C with shaking, and OD measurements were taken every half an hour. OD measurements were recalculated into biomass and relative biomass. Based on experimental data, the model parameters were fitted.

### Experimental long starvation scenario

#### Preparation of experimental populations

For the long starvation scenario, cells stored at -70°C were thawed, merged accordingly to type (Q, NQ) and washed in sterile water 3 times. Q and NQ cells were diluted to equal density (OD = 0.8, all OD measurements were taken with Multi-Mode Microplate Reader SpectraMax iD3, with λ = 600 nm) in sterile water. Then six types of experimental cultures (Q monoculture, NQ monocultures and Q+NQ mixed cultures, each in sterile H_2_O and spent medium) were set up, each type of experimental populations were prepared in 5 repetitions, 5 ml each, giving all together 30 independently starved cultures. Mixed cultures were set up by mixing Q and NQ cells in 3:1 v/v proportions. Mixed cultures are treated as reference point because they mimic naturally occurring Q/NQ balance in the S288C yeast strain. Two **starvation media** were used: sterile water (*simple environment*) and spent medium (*complex environment*) (Fig.1B). To harvest spent medium, yeast cells of the same prototrophic S288C strain were inoculated in fresh YPD for 4 days, then cells were pelleted and supernatant was filtered and placed in a sterile container. The remaining cells were discarded. Lack of viable cells in the spent medium was confirmed by spreading samples of the harvested medium on YPD plate and incubation for 5 days in 30°C. No colony growth on these plates was observed.

For the long starvation scenario the cultures were kept at 30°C with shaking for 6 weeks. Samples from starving cultures were weekly checked for regrowth ability (“*regrowth procedure*”), starting 1 week (7 days) after the start of the experiment.

#### Regrowth procedure

From each starving culture, a 275 µl sample was taken and spun down, supernatant (starvation medium) was discarded, and cells were resuspended in 550 µl of fresh YPD medium. Then 200 µl was placed in a 96-well plate (flat bottom) in two repetitions. Additionally, *fresh cells* (inoculum: cells of the same S288C strain from liquid YPD medium incubated o/n in 30°C) were placed into the plate as a control. The plate was covered with transparent incubation foil and placed in the reader (SpectraMax iD3 Multi-Mode Microplate Reader) for 70h at 30°C with shaking. OD measurements (λ = 600 nm) were taken every 30 minutes. The procedure was repeated weekly through the 6 weeks of the starvation experiment (Fig 1C).

### Experimental short starvation scenario

Q and NQ cells were acquired in the same way as described above. Then, immediately after fractionation, Q and NQ cells were diluted to equal density (OD = 0.4), and the mixed culture was prepared by mixing Q and NQ cells in 3:1 proportion. Q monoculture, NQ monoculture and mixed culture were suspended in fresh liquid YPD medium. Then 150 µl were put in a flat bottom 96-well plate in 16 repetitions for each culture type. The plate was covered in transparent incubation foil, placed in the reader for incubation at 30°C with shaking for 24 hours. OD measurements (λ = 600 nm) were taken every 30 minutes.

### Relative biomass analysis

OD values were first converted into biomass [cell number] according to the equation:

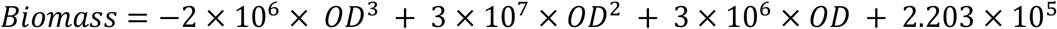

The biomass values given throughout the manuscript are the number of cells present in 200µl, which is the total volume in a well (in 96-well plate) during regrowth. The equation was established by combining OD measurements (λ = 600 nm) and cell counting in a flow cytometer (Beckman Coulter CytoFLEX) after staining with propidium iodide (PI).

The data analysis was conducted after 24 hours (out of 70 hours) of regrowth, as this is the best time frame to capture differences between experimental populations. We compared how monocultures regrow after starvation in comparison to mixed cultures by relative biomass analysis. Relative biomass of a given monoculture was calculated as the ratio of its biomass and the average biomass of mixed cultures at a given time point of regrowth (“*weekly procedure*”). Relative biomass of mixed culture equals 1 on average (Fig. 2, Fig. 3).

**Fig. 2.**
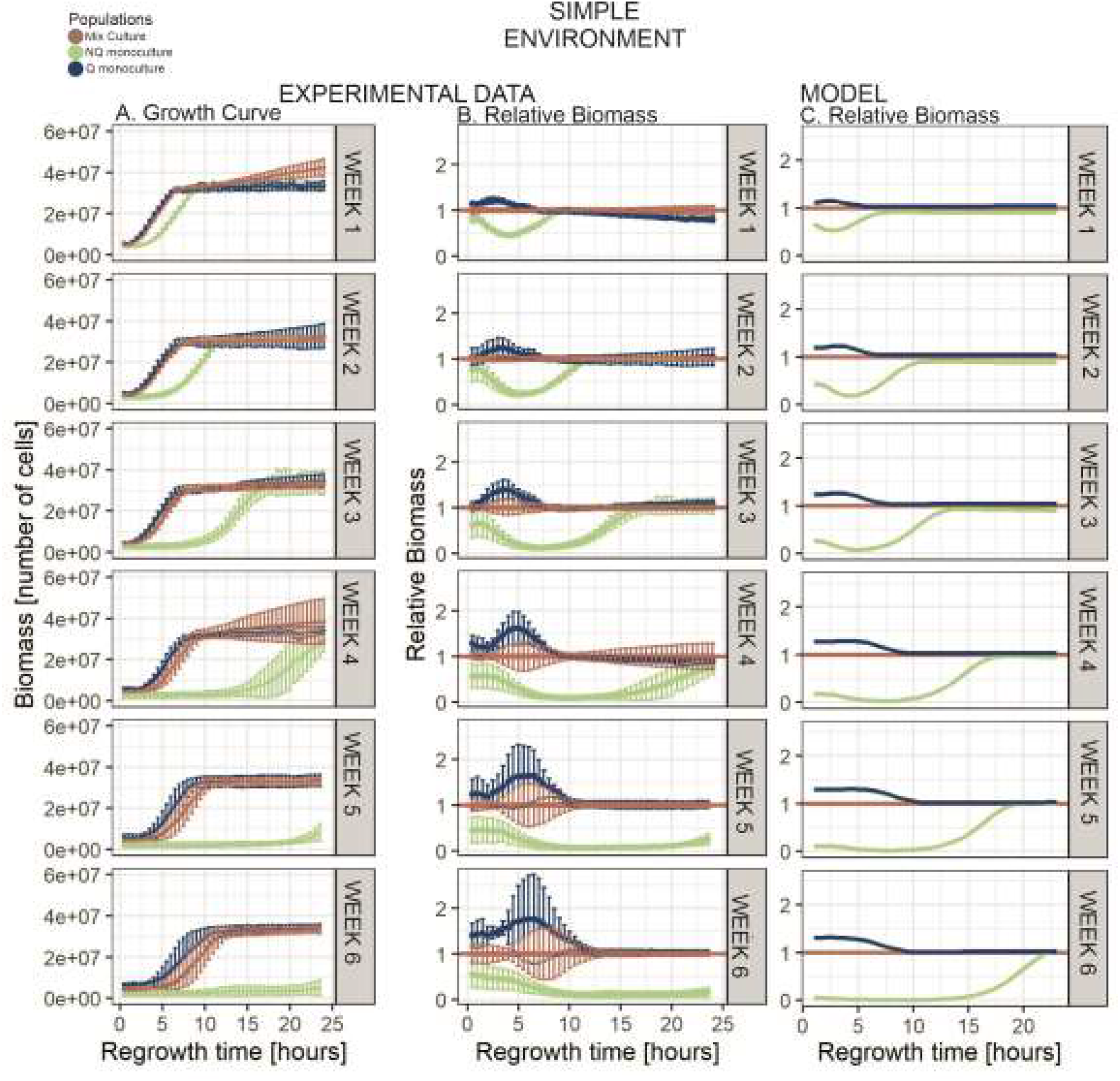
Simple Environment results Results from the starvation in the simple environment. Subsequent vertical graphs illustrate the length of starvation in weeks (1 to 6). The regrowth time (in hours) illustrate time elapsed since placing samples from starved experimental cultures into fresh media for regrowth. Points illustrate the averaged results from five repetitions (except of week 5 and 3 for NQ monoculture where there were four repetitions each), and error bars show standard deviation. The Q (navy) and mix (brown) cultures display similar growth curves until 3^rd^ week of starvation. When starved for 4 weeks or longer, growth curve of Q monoculture is increasing sharper, reaching flat shape of stationary phase earlier. Growth curve of the NQ monoculture (green) since 1^st^ week of starvation is the flattest and it reaches stationary phase later than both mix and Q cultures.

**Fig. 3.**
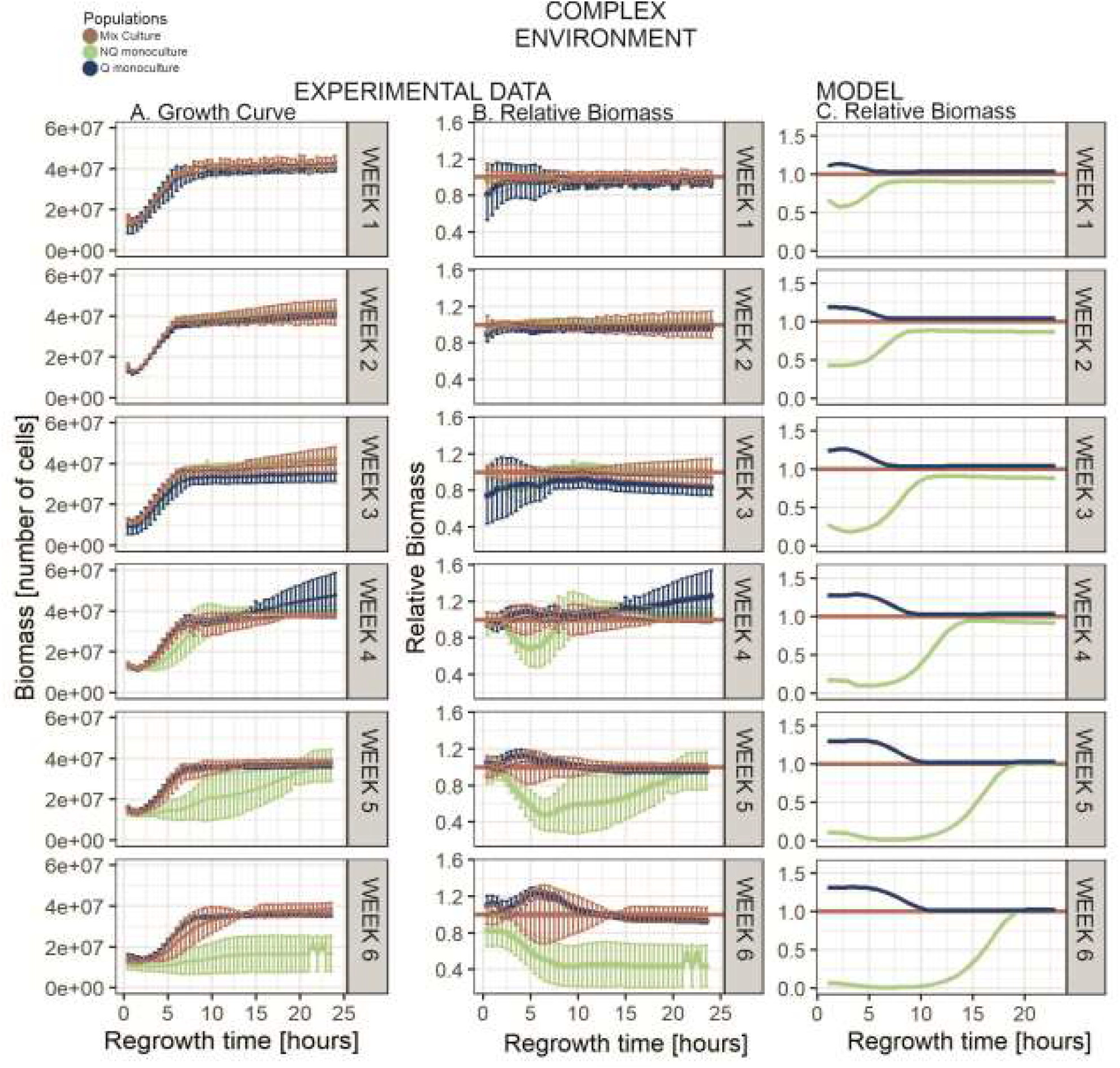
Complex environment results Results from starvation in the complex environment. Subsequent vertical graphs illustrate the length of starvation in weeks (1 to 6). The regrowth time (in hours) illustrate time elapsed since placing samples from starved experimental cultures into fresh media for regrowth. Points illustrate the averaged results from five repetitions (except of week 6 for NQ monoculture where there were three repetitions), and error bars show standard deviation. The growth curves of all starved populations are similar until 3^rd^ week of starvation. When starved for 4 weeks or longer, the growth curve of NQ monocultures gradually flatten. Q and mixed cultures display similar shapes of growth curves throughout whole starvation time.

To compare the effect of the starvation medium on the experimental populations survival (Fig. 4), relative biomass was calculated as the ratio of the biomass of an experimental culture of a given cell type starved in the complex environment (spent medium) and the biomass of an experimental culture of the same cell type starved in the simple environment (sterile water), at a given time point.

**Fig. 4.**
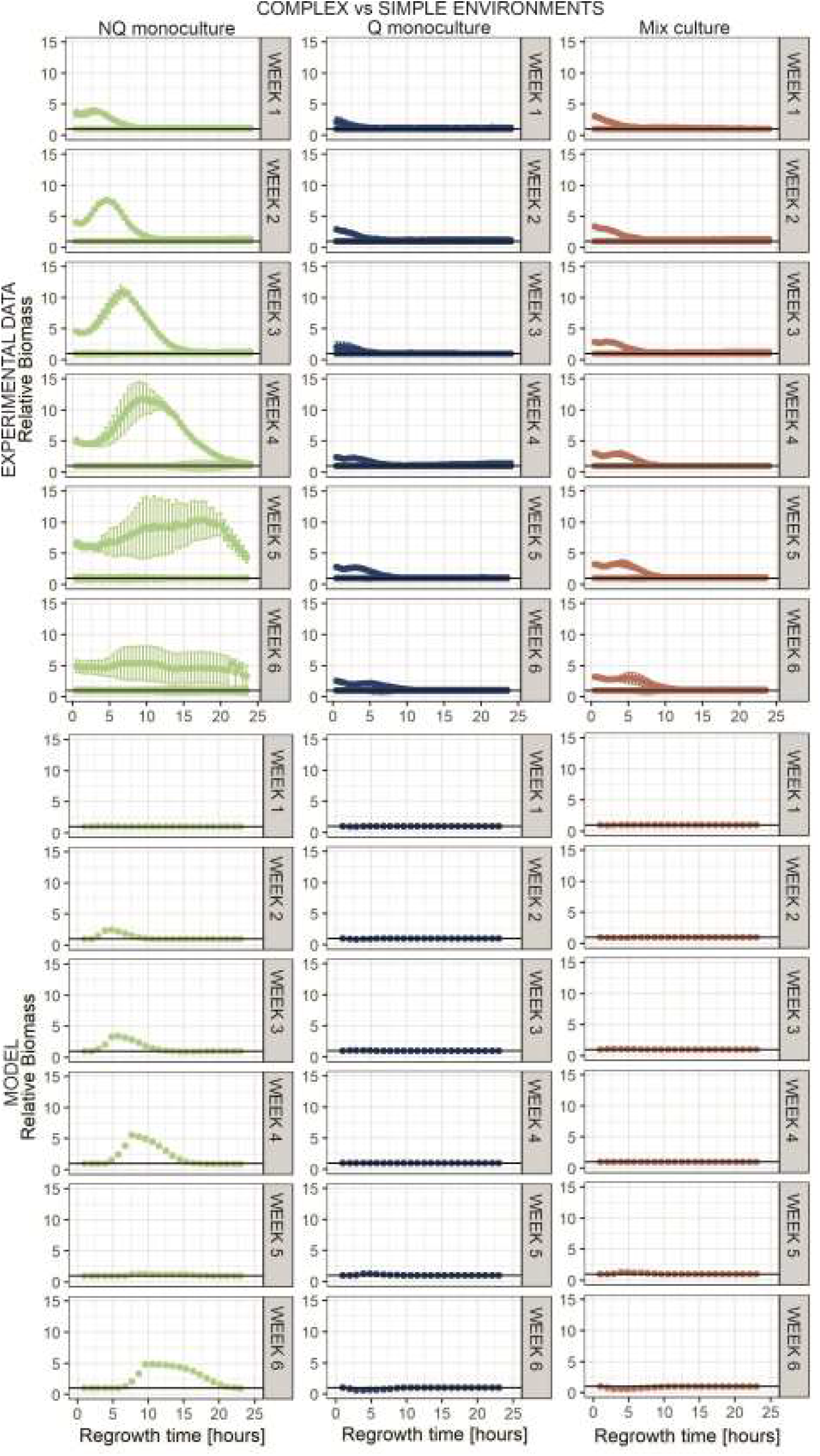
Complex vs simple environment Starvation environments’ impact on cultures’ starvation survival. Subsequent vertical graphs illustrate length of starvation in weeks (1 to 6). Regrowth time (in hours) illustrate time elapsed since placing samples from starved experimental cultures into fresh media for regrowth. Relative biomass was calculated as a ratio of biomass of given cell type culture in complex environment to the same cell type culture biomass in simple environment (dots). Biomass of a culture in simple environment equals 1 (solid line). NQ monocultures survive relatively better when starved in complex than simple environment. Q and mix cultures benefit slightly at the beginning of regrowth (up to 10 hours) when starved in complex environment.

Statistical analysis of several chosen timepoints were conducted via ANOVA test as biomass ∼ experimental culture, followed by post hoc Tukey tests.

### Lag phase length analysis

Lag phase length was defined as the time needed for a population to increase its OD by 0.01 from the OD at the beginning of regrowth. Lag phase lengths of different cultures after a given week of starvation were compared using an two-sided T-test (S2 Table1, S3 Table2).

### Model description

The model simulations mirror the experimental procedures in which we first starve the cultures and then let them regrow in fresh media. The model represents the biological reality that differentiation into Q and NQ cells starts when the nutrients are nearly depleted, and that this differentiation is not instantaneous (S4 Fig.1, S5 Fig.2). The model tracks the concentration of limiting resources, nutrients available for recycling and various types of cells over time. Population dynamics in the starvation phase is determined by the death rates of Q and NQ (calculated based on [9], and nutrient recycling (which is assumed to occur in complex, but not in simple environments). The population dynamics during regrowth on fresh media is based on the standard Michaelis-Menten kinetics, taking into account different lengths of the lag phase for Q and NQ cells.

The model is based on previous ODE models that use a bottom-up approach to track the population dynamics in specific ecological contexts [27–29]. The detailed description of the model and model parameters can be found in the Supplementary materials (S1). The numerical solutions of the model were obtained using Matlab 2018a, and the parameters were fitted using the R global optimization package DEoptim [30].

## Results

### I. Long starvation in simple environment

#### Experimental data: Advantage of Q monocultures is dynamic

Higher proportions of stress-resistant Q cells should provide better population survival during starvation, which can be measured as exponential biomass increase after lag phase in a fresh growth medium. Accordingly, Q monocultures were expected to synchronously restart divisions soon after nutrient restoration and reach stationary phase density before other cultures. Indeed, at the beginning of the regrowth procedure, after first week of starvation, Q monocultures have a biomass advantage over mixed cultures (after 2 hours of regrowth, average biomass of Q cultures = 8.22×10^6^ cells and average biomass of mix cultures = 6.85×10^6^ cells; p = 0.0005, Fig. 2, S10.Fig7). Yet, the advantage of Q monocultures is small, and during further weeks of experiments, it is not significant (after 2 hours of regrowth: 2^nd^ week: p = 0.33, 3^rd^ week: p = 0.129, 4^th^ week: p = 0.177, 5^th^ week: p = 0.404) except for 6^th^ week (p = 0.037).

#### Experimental data: NQ monocultures regrow more slowly

The increase in biomass of NQ monocultures is slower than that of the other experimental populations (Fig. 2). The significant disadvantage is already visible after 1^st^ week of starvation, where initially NQ monocultures have a lower biomass than other cultures (after 2 hours regrowth: average biomass of NQ cultures = 4.39×10^6^cells; post-hoc Tukey test: NQ-Q: p = 1.31×10^−8^, NQ-mix: p = 1.74×10^−6^), and only reach the biomass of Q cultures and mixed cultures after 10 hours of regrowth (Fig. 2, 10 hours regrowth: average biomass of Q cultures = 3.27×10^7^cells, biomass of mixed cultures = 3.32×10^7^cells and biomass of NQ cultures = 3.29×10^7^cells; p > 0.05 for both NQ-Q and NQ-mixed comparisons). Also the lag phase of NQ monocultures is longer than those of other experimental populations (Fig. 5, S2.Table1) The biomass differences between NQ and other cultures increases with starvation time. During further weeks of starvation, NQ monocultures need more and more time to reach stationary phase density, exceeding 24 hours after the 4^th^ week of starvation (Fig. 2).

**Fig. 5.**
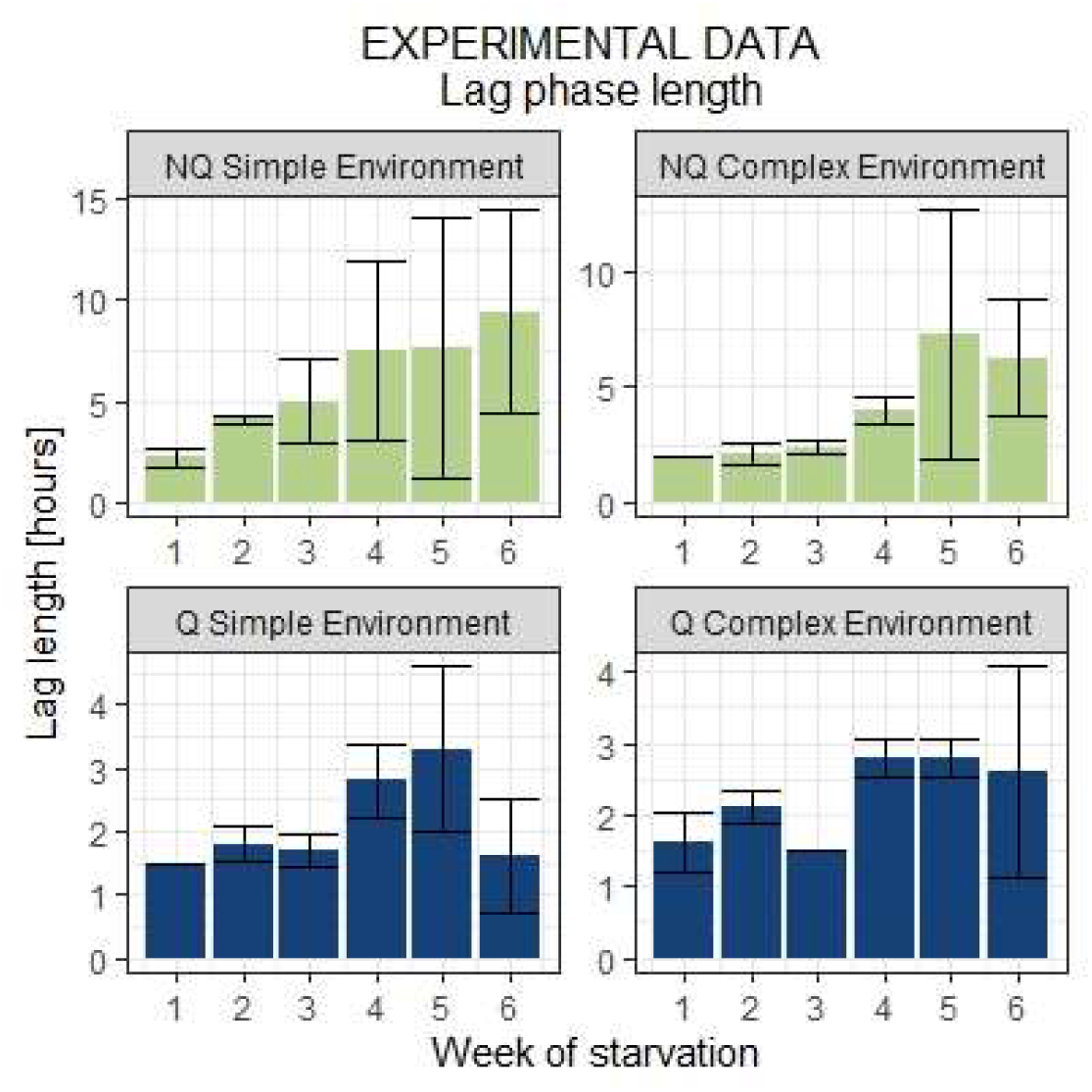
Average lag phase length of experimental monocultures during starvation. Bars represent standard deviation. Length of lag phase evidently increase for NQ monoculture and it is especially visible in simple environment. Meanwhile, for Q monoculture lag phase length is almost constant through whole starvation time.

We compared lag phase lengths for Q and NQ monocultures starved in simple environments. The length of the lag phase increases with starvation time (p = 3.58×10^−8^, Fig. 5, S2 Table1, S3 Table2, S9.Fig.6). While at the beginning of the experiment the lag phase is on average 1.5 hours for Q and 2.2 hours for NQ monocultures, the lag phase of NQ monocultures increases to an average 9.4 hours after 6^th^ week of starvation. The increase of the lag phase of Q monocultures is much lower than that of NQ (p = 0.0003, S2 Table1, S3 Table2) and its length is on average 1.6 hours after 6 weeks of starvation.

### II. Analysis of model predictions

#### Model

In order to shed light on possible reasons for the advantage of Q monocultures over mixed cultures, and on how this advantage depends on the length of starvation and regrowth time, we used a mathematical model that tracks the ratio of Q and NQ cells over time, as well as the concentrations of limiting nutrients (see S1 file for a detailed description of the model). The model reproduced the experimental results from the simple starvation environment: (i) the advantage of Q monocultures over other cultures increases with the starvation time and (ii) Q’s advantage is noticeable at the beginning of regrowth but with increasing length of regrowth, this advantage decreases and finally disappears (Fig. 2C).

The model suggests that there are two factors that drive those results: death rate during starvation and lag length (if they were equal for Q and NQ cells both cell types would follow the same starvation and regrowth dynamics, (S6)). In particular the dependence of the relative biomass of different culture types on the regrowth time results from the fact that Q cells have a shorter lag length than NQ cells (S7.Fig.4A). In contrast, the dependence of the relative biomass on the starvation time can result from either (i) higher death rate for NQ cells than Q cells during starvation (S7. Fig4B) or (ii) lag lengths consistently growing with the starvation time (S7. Fig4C).

Based on modelling, we also predicted that nutrient recycling is a potentially important factor influencing survival during starvation. The model suggests that NQ cells should be the ones that benefit from more complex starvation environment. Nutrient reusability helps NQ monocultures regardless of their lag time: the model yields analogous results even if both Q and NQ cells have no lags during regrowth (S7 Fig.4 D). This suggests that it is the nutrient recycling that drives the difference in NQ survival in the two environments.

### III. Long starvation in complex environment

#### Experimental data: Nutrient recycling is crucial for NQ cells’ survival

To test the model predictions we repeated our long starvation experiment in the complex environment, where nutrient recycling is possible. Experimental results revealed that there is no significant difference between populations at the beginning of regrowth up to 5^th^ week of starvation (after 2 hours regrowth, p > 0.05 for all NQ-Q, NQ-mixed and Q-mixed comparisons, (S10 Fig.7). Up to the 2^nd^ week of starvation, NQ monocultures regrowth is similar to Q and mixed cultures regrowth (Fig. 3A, 3B), reaching the same biomass after sufficient regrowth time (maximal density after ∼5 hours, 1^st^ week, on average: biomass of Q cultures = 3.04×10^7^cells, biomass of NQ cultures = 3.23×10^7^cells, biomass of mix cultures = 3.21 × 10^7^cells; p > 0.05 for all NQ-Q, NQ-mix and Q-mix comparisons). After further weeks of starvation, the differences in regrowth efficiency between NQ monocultures and mixed cultures on the one hand, and Q monocultures and mixed cultures on the other hand, gradually increase. After the 4^th^ week of starvation, NQ monocultures reach stationary phase density within 10 hours of regrowth and a week later, they need almost 24 hours to reach the same biomass (Fig 3). In terms of length of regrowth, the same pattern can be observed as described previously – with increasing regrowth time, differences between populations progressively decrease and finally disappear when regrowth time is long enough for all cultures to reach stationary phase density (Fig. 3, S8. Fig.5). Direct comparison of the simple and complex starvation environment revealed that the biomass of NQ monocultures can be even 11 times higher (4^th^ week of starvation, regrowth time from 8.5 to 11.5 hours) when starved in the complex environment (Fig. 4) than when starved in the simple environment. The model results follow the general pattern of experiments (except for NQ monocultures in 5^th^ week – probably due to an unusually long experimental lag phase in the complex environment (S3 Table 2), although some quantitative differences may also be caused by large variance in experimental data. Regrowth abilities of Q monocultures and mixed cultures were influenced by starvation medium to a lesser extent (Fig. 4).

### IV. Short starvation scenario

#### NQ monoculture have biomass advantage in short starvation

Since the disadvantage of NQ is smaller when starvation time is short, we used the short starvation experiment to verify if NQ cells can have an advantage over Q cells. Our model predicted that this could be the case if the lag of NQ cells is shorter than that of Q cells (Fig. 6). To test this experimentally, we isolated Q and NQ cells from cultures that had been in stationary phase for 4 days (short starvation). Q and NQ monocultures as well as mixed cultures were prepared and the cultures were placed into a fresh rich medium for regrowth. After such short starvation the NQ monocultures restarted growth faster than the Q and mixed cultures (average lag length for Q monoculture = 2 ± 0,0 hours and for NQ monoculture = 1 ± 0,0 hour, p = 2.2×10^−16^). Relative biomass analysis revealed that the advantage of NQ persists up to 8 hours after inoculation (Fig. 6, NQ-mix: p = 8.08×10^−4^, NQ-Q: p = 5.5×10^−7^).

**Fig. 6.**
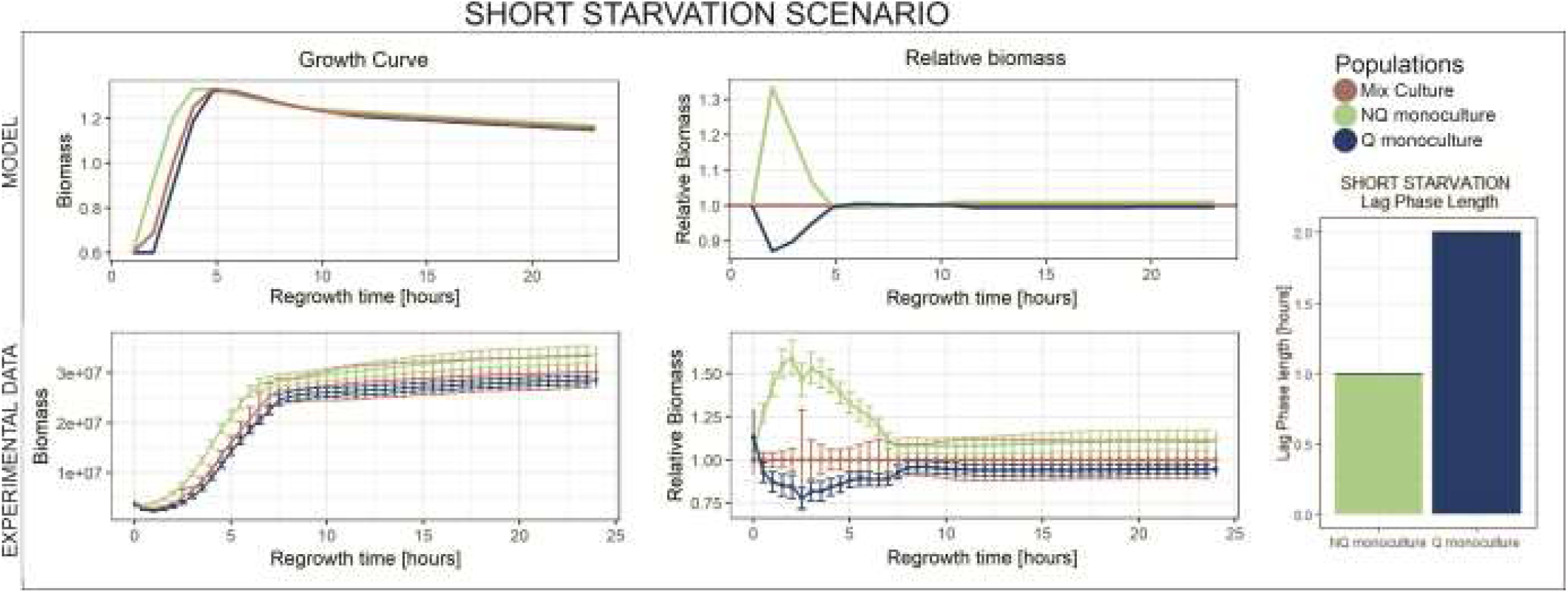
Short starvation scenario We modeled experimental populations in short starvation scenario. NQ monoculture gain advantage over both Q and mix cultures up to 5 hours of growth. The model suggested that this advantage is due to shorter lag phase of NQ cells when starved for short period of time. The model results were confirmed by laboratory experiment. When starved only for 4 days, NQ monoculture indeed outcompete mix and Q cultures at the beginning of regrowth (up to ∼7 hours). Moreover, the lag phase of NQ cells is shorter than Q cells (the bars represent average from 16 repetitions).

## Discussion

Here we examined six types of experimental populations – three cell composition types (Q and NQ monocultures, and Q+NQ mixed cultures) in two starvation media (sterile water and spent medium, representing simple and complex environments, respectively). Populations were starved for 6 weeks, and regrowth abilities (biomass increase) and lag phase length in fresh rich medium were monitored after each week. We also conducted an experiment testing experimental populations in a very short, 4 day starvation scenario.

### Q cells are adapted to survive long starvation

We showed that Q monocultures have a clear biomass advantage over NQ monocultures after long starvation. The difference is especially pronounced when regrowth times are short (up to 10 hours of growth), and Q’s advantage increases when populations are starved longer (Fig. 2, Fig. 3). The biomass of Q monocultures is also higher on average than those of mixed cultures (Fig. 2, Fig. 3), although the difference doesn’t appear to be statistically significant. It can be explained by the fact that mixed cultures are composed of 75% of cells in the quiescent state. The advantage of Q over mixed cultures was also confirmed by the model (Fig. 2C, Fig. 3C). We can therefore hypothesize that it is mostly quiescent cells that are responsible for the survival and regrowth abilities of mixed cultures. This is a consequence of Q cells dying at a lower rate than NQ cells under starvation [9,11] and having shorter lags when entering regrowth in rich medium [11]. However, this advantage decreases and finally disappears when regrowth times increase, because after a sufficiently long time for regrowth all populations reach their carrying capacities regardless of the initial size and lags (Fig. 2, Fig. 3, S8.Fig5). This is because the maximal biomass in any case is determined by nutrient abundance in a given medium. Thus, when the regrowth time is long enough, any biomass advantage would gradually decrease and finally disappear.

### Nutrient recycling increase survival of NQ cells during long starvation in complex environment

We showed that while the starvation medium has little or no impact on Q monocultures and mixed cultures, the NQ monocultures survive relatively better in complex environment (Fig. 4). Both the simple and the complex starvation media were lacking glucose, however, in the complex environment, which consists of spent medium, nutrient and metabolite recycling was possible. It is because some nutrients, such as some amino acids, remain in the spent medium even after the onset of growth and starvation, and because additional nutrients may be released from dead cells during long starvation [27]. In addition, it had been demonstrated that excretion of nutrients and metabolites into the environment is a natural property of yeast populations and that cells can cooperatively exchange exometabolites [27,31,32]. In simple environment the nutrients released from dead cells are unlikely to be reused during starvation because the environment is too poor to provide all types of nutrients necessary for growth. However, in complex environment, released nutrients can cover missing compounds and together with nutrients remaining in spent medium, enable increased survival. Indeed, death and nutrient recycling has been demonstrated as potentially crucial in bacterial communities [27]. We captured this phenomenon in our model by varying to what extent nutrients can be recycled. Experiments performed in complex starvation media confirmed model predictions that NQ monocultures do better when nutrient recycling is possible (Fig. 3, 4). This could be because the nutrient recycling simply helps to reduce the effect of death rates being higher for NQ than Q cells.

### NQ outperform Q cells when starvation is short

Shorter starvation times result in smaller growth advantages of Q monocultures compared to mixed cultures (Fig. 2, 3). This raises the question of whether there could be scenarios in which entering quiescence is not beneficial at all. The model suggested that this could be the case if the lag phase for NQ cells was shorter than for Q cells (Fig 6). Indeed we showed that in case of very short starvation (4 days), lag phase length is shorter for NQ cells (Fig.6). In particular, we showed that when a culture faces a very short starvation period, cells that have switched to the Q state experience longer lags than those that remained in the NQ state (Fig. 6). As a consequence, at the beginning of regrowth after short starvation, NQ monoculture gains an advantage over Q and mixed cultures.

### Presence of both Q and NQ cells is adaptive for population under specific ecological scenarios

Multiple studies have demonstrated advantages of quiescent cells over any other phenotype when populations are facing stressful conditions. Simple environments with a single, highly stressful factor (such as heat shock or toxins) indeed favour more resistant quiescent cells [9,11]. Such tests provide important information, however, natural populations usually face fluctuating environmental changes, often with gradually increasing stress [24,33].

Overall, our results show that switching to the Q state may not always be adaptive, and that the benefits of this physiological transition depends on the ecological context. When the starvation period is very short (e.g. 4 days) and beneficial conditions are restored, cells that do not switch to quiescence benefit from a shorter lag phase. On the other hand, quiescent cells survive long starvation much better. Moreover, when nutrient recycling during starvation is possible, NQ cells perform as well as Q if the starvation is no longer than 2 weeks.

In particular, the fact that natural populations are phenotypically heterogeneous during starvation and are composed of both Q and NQ cells may not be a side effect of an imperfect switch to quiescence, but may actually be a proper bet hedging strategy under uncertain ecological conditions. It will be interesting to further test these assertions using evolution experiments in which populations are subject to different lengths and frequencies of periods of starvation and regrowth.

## Data availability

Experimental data generated in laboratory during this study will be available upon request or deposited to online server if the ms is accepted for publication.

## Code availability

The model code is publicly available on GitHub. https://github.com/bognabognabogna/Q-NQ-data-analysis/tree/master/Matlab_scripts

## Acknowledgements

We would like to thank Donna Katie McCullough, Klaudia Wilk and Anna Smoter for help in laboratory and Katarzyna Tomala, Richard Lindsay and Rafael Carlos Reding for insightful comments. The work was funded by the National Science Centre, Poland via an OPUS grant to D.W-S. No. 2017/25/B/NZ8/01035, the Biology Department Research subsidy No. N18/DBS/000003 and N18/MNW/000013 to MO. MD was supported by NSERC Award Number: 041931Y. BS was supported by Polish National Agency for Academic Exchange by grant No PPN/PPO/2018/00021/U/00001.

